# The *Drosophila melanogaster* ortholog of RFWD3 functions independently of RAD51 during DNA repair

**DOI:** 10.1101/844779

**Authors:** Juan Carvajal-Garcia, Evan R. Gales, Dale A. Ramsden, Jeff Sekelsky

## Abstract

Repair of damaged DNA is required for the viability of all organisms. Studies in *Drosophila melanogaster*, driven by the power of genetic screens, pioneered the discovery and characterization of many genes and pathways involved in DNA repair in animals. However, fewer than half of the alleles identified in these screens have been mapped to a specific gene, leaving a potential for new discoveries in this field. Here we show that the previously uncharacterized mutagen sensitive gene *mus302* codes for the *Drosophila melanogaster* ortholog of the E3 ubiquitin ligase RING finger and WD domain protein 3 (RFWD3). In human cells, RFWD3 promotes ubiquitylation of RPA and RAD51 to facilitate repair of collapsed replication forks and double strand breaks through homologous recombination. Despite the high similarity in sequence to the human ortholog, our evidence fails to support a role for Mus302 in the repair of these types of damage. Last, we observe that the N-terminal third of RFWD3 is only present in mammals and absent in the rest of vertebrates and invertebrates. We propose that the additional N-terminal portion accounts for the acquisition of a new biological function in mammals that explains the functional differences between the human and the fly orthologs, and that *Drosophila* Mus302 may retain the ancestral function of the protein.

## Introduction

DNA damage repair consists on a processes that detect and fix changes in the DNA molecules of cells; DNA repair is required for cell and organismal viability. *Drosophila melanogaster* has been an important model in the discovery of genes involved in DNA damage repair (Sekelsky, 2017). In the 1980s and 1990s, dozens of mutants hypersensitive to the DNA alkylating agent methyl methanesulfonate (MMS) were isolated (mutagen-sensitive genes, *mus*) (Boyd, Golino, Shaw, Osgood, & Green, 1981; Laurencon et al., 2004; Mason, Green, Shaw, & Boyd, 1981). Mapping and characterization of these mutants has led to important insights into DNA repair mechanisms not only in fruit flies but also in humans (Andersen et al., 2009; Chan, Yu, & McVey, 2010). However, the majority of these mutations have yet to be characterized (*e.g.*, 20 of 27 on chromosome 3), providing a useful resource to continue improving our understanding of DNA repair. In this study, we map one these uncharacterized complementation groups, *mus302*, and show that the gene encodes the ortholog of the human RING finger and WD domain protein 3 (RFWD3).

RFDW3 is an E3 ubiquitin ligase that targets the single-stranded DNA binding protein Replication Protein A (RPA) (Elia et al., 2015; Liu et al., 2011), the recombinase RAD51 (Inano et al., 2017) and the tumor suppressor p53 (Fu et al., 2010) after DNA damage in humans. The fate of the ubiquitylated proteins is not clear, as different groups report different conclusions (Elia et al., 2015; Inano et al., 2017). In humans, RFWD3 has been shown to be involved in the restart of hydroxyurea (HU)-stalled replication forks, the repair of Tus/ter collapsed forks through homologous recombination (HR), as well as repair of *I-Sce*I-mediated double strand breaks (DSBs) (Elia et al., 2015). Human cells deficient in RFWD3 are also hypersensitive to the DNA crosslinking agent mitomycin C (MMC), ionizing radiation (IR), and HU (Feeney et al., 2017; Inano et al., 2017). *RFWD3* mutant cells exhibit increased foci of RPA and RAD51 when treated with MMC (Feeney et al., 2017). Consistent with these observations, RFWD3 localizes to replication forks in a proliferating cell nuclear antigen (PCNA)-dependent manner (Lin et al., 2018). In addition, RFWD3 is phosphorylated by the DNA damage response kinase ATR (and possibly ATM) (Feeney et al., 2017; Fu et al., 2010), and this may be required for its function. Finally, patients biallelic for inactivating mutations in *RFWD3* display Fanconi Anemia-like symptoms, so this gene has also been named *FANCW* (Knies et al., 2017).

Here we show that flies with mutations in *mus302* display no hypersensitivity to HU or IR, suggesting that Mus302 is not involved in the repair of collapsed replication forks or DSBs, despite its orthology to RFWD3. Moreover, these flies have no apparent defects in a gap repair assay of synthesis-dependent strand annealing (SDSA), one of the most common pathways for homologous repair of DSBs. We also provide evidence that Mus302 acts independently of the *Drosophila* ortholog of RAD51 (Spn-A) in repair of DNA damage caused by MMS. Last, we observe that two known ATR phosphorylation sites in human RFWD3 are missing in Mus302, consistent with a role of this protein in DNA repair outside of S phase. Taken together, our findings show that the *Drosophila* ortholog of RFWD3 functions differently from the human one, suggesting it may be used to reveal new roles of the protein in humans.

## Materials and methods

### Drosophila stocks

Drosophila stocks were kept at 25°C on standard cornmeal medium. Flies with mutant *mus302* alleles were obtained from the Bloomington Drosophila Stock Center (BDSC) and are described in (Boyd et al., 1981) and (Laurençon et al., 2004) (*mus302*^*D1*^, *mus302*^*D2*^, *mus302*^*D3*^, *mus302*^*Z1882*^, *mus302*^*Z4933*^ and *mus302*^*Z6004*^). To generate a wild type *CG13025* transgene, the coding sequence plus the intron of this gene was amplified with 1187 bp upstream of the ATG and 271 bp downstream of the stop codon and cloned into a plasmid containing an *attB* site and a *w*^+^ gene. The plasmid was injected into the Bloomington stock number 9738 (*y*^*1*^ *w*^*1118*^; *PBac*{*y*^*+*^*-attP-9A*}(*VK00020*) (Genetivision) and two independent isolates (A and B) were generated. The *3L* deficiency stocks *Df(3L)ED4606* (deletes 16,087,484-16,780,123) and *Df(3L)ED4674* (deletes 16,661,284-17,049,418) were obtained from BDSC (stock numbers 8078 and 8098). *spn-A*^*057*^ and *spn-A*^*093A*^ mutations are described in (Staeva-Vieira, Yoo, & Lehmann, 2003).

### DNA damage sensitivity assays

Sensitivity to DNA damaging agents was assessed as in (Holsclaw & Sekelsky, 2017; Sekelsky, 2017). Three males and five females heterozygous for the indicated mutations were crossed and allowed to lay eggs for three days (untreated brood). They were then transferred into a new vial and allowed to lay eggs for two days (treated brood). The treated brood was exposed to the indicated dose of methyl methanesulfonate, hydroxyurea or ionizing radiation (source: ^137^Cs). The fraction of homozygous mutants for both broods was calculated per vial. Survival was calculated as the fraction of homozygous mutants in the treated brood over the fraction of homozygous mutant in the untreated brood.

### Allele amplification and sequencing

For sequencing the coding region of *CG13025* in flies with mutant *mus302* alleles, each allele was crossed to the deficiency line *Df(3L)ED4606* and DNA was extracted from a male as in Adams *et al.* (2003). *CG13025* was amplified with the high-fidelity polymerase PrimeSTAR HS (Takara) and sequenced by Sanger sequencing (Eton). Sequences from the mutant alleles were compared to the presumed original wild-type alleles in flies from the corresponding screen by sequencing *CG13025* from the *mus312*^*D1*^ and *mus312*^*Z1973*^, which were isolated in the same screens. Allele-specific PCRs were developed for the *mus302*^*D1*^ and the *mus302*^*Z1882*^ mutations to generate recombinants.

### Gap repair assay

The *P*{*w*^*a*^} gap repair assay was performed as a slightly modified version of the one described by Adams *et al.* (2003). In short, females containing the *P*{*w*^*a*^} element and heterozygous for the *mus302*^*D1*^ allele were crossed to males carrying *P* transposase and heterozygous for the *mus302*^*1882*^ allele. Single male progeny of this cross expressing *P* transposase and either heterozygous for *mus302*^*Z1882*^ or heteroallelic for both *mus302* mutations were crossed to females with the compound *X* chromosome *C(1)DX*. Male progeny that did not inherit transposase were scored as “red-eyed” (SDSA), “white-eyed” (alt-EJ), or “apricot-eyed” (mostly no excision but possibly full restoration of *P*{*w*^*a*^}).

### Sequence alignment

The sequences for the RFWD3 orthologs in *Homo sapiens, Mus musculus, Gallus gallus, Xenopus tropicalis, Danio rerio, Strongylocentrotus purpuratus*, and *Drosophila melanogaster* were downloaded from Ensembl. Protein sequences were aligned in ClustalX 2.1 (Larkin et al., 2007) and edited in GeneDoc 2.7.000 (Nicholas et al., 1997).

### Statistical analysis

Statistical analyses were performed with Prism 8 (GraphPad). Tests are indicated in figure legends. Statistical significance is defined as *p*<0.05.

### Data and reagent availability

*Drosophila* stocks, plasmids, and primer sequences are available upon request. Supplemental figures S1 and S2 have been uploaded to FigShare.

## Results and discussion

### *mus302* encodes the *Drosophila melanogaster* ortholog of the human RFWD3

We sought to map one of the uncharacterized mutagen-sensitive (*mus*) complementation groups in the third chromosome of *Drosophila melanogaster* to a defined chromosomal location. 20 of the 27 groups are yet to be mapped so we focused on *mus302. mus302* alleles (*D1* through *D6*) were first isolated by Boyd *et al.* (1981) as conferring hypersensitivity to methyl methanesulfonate (MMS). Laurençon *et al.* (2004) found five additional *mus302* mutations in another screen (*Z1882, Z4933, Z6004, Z2530* and *Z5541*). We confirmed that Boyd’s *mus302* complementation group corresponded to Laurençon’s by testing the sensitivity to 0.025% MMS in *mus302*^*D1*^/*mus302*^*Z1882*^ heteroallelic mutants and observing that this dose is lethal to these mutants but not their heterozygous siblings (Fig. 1a).

**Figure 1.**
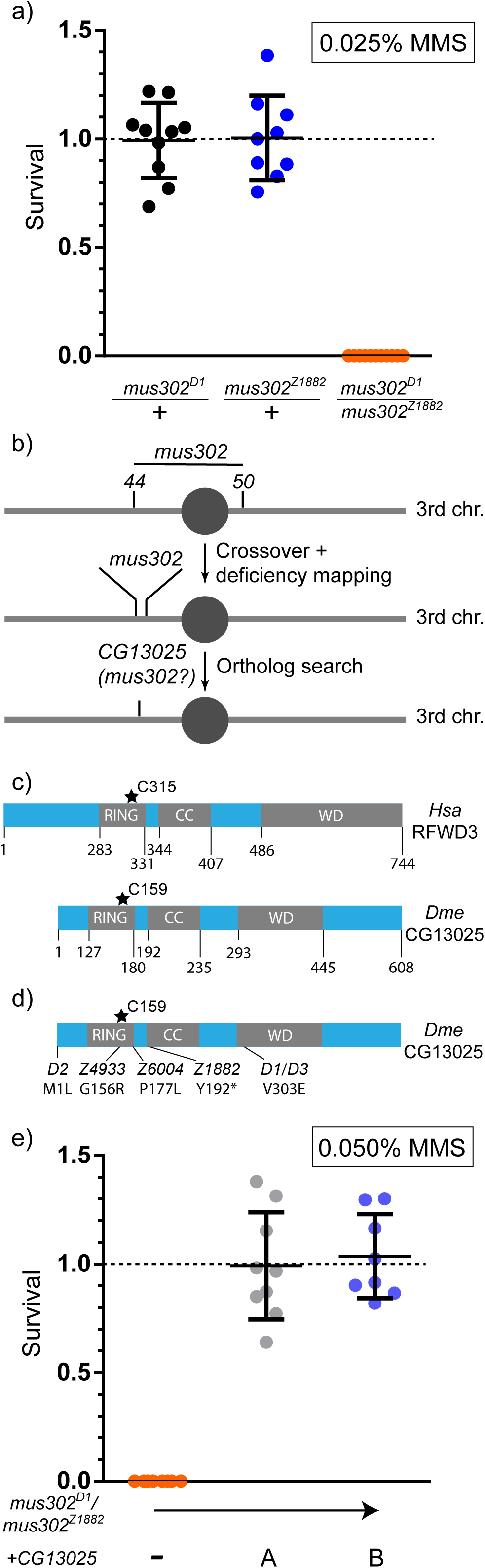
Mus302 is an ortholog of RFWD3. **A)** Survival of flies exposed to 0.025% methyl methanesulfonate of the indicated genotype with respect to the untreated progeny from the same parents. Chromosomes with wild-type *mus302* had the *mus312*^*Z1973*^ mutation (crossed to *mus302*^*D1*^) or the *mus312*^*D1*^ mutation (crossed to *mus302*^*Z1882*^). Horizontal dashed line at Y=1 indicates 100% survival. **B)** Schematic of the third chromosome of *Drosophila melanogaster* (circle represents the centromere, not to scale). Numbers represent the genetic position of *st* (44) and *cu* (50). After crossover mapping, we observed that *mus302* was close to *st*. Deficiency mapping narrowed the region 22 possible genes. The predicted gene *CG13025* was our primary candidate. **C)** Schematic of *Homo sapiens* (*Hsa*) RFDW3 and *Drosophila melanogaster* (*Dme*) CG13025. RING finger and WD domain boundaries were determined with the Conserved Domain tool from NCBI (Marchler-Bauer et al., 2013) and the coiled-coil motif with DeepCoil (Ludwiczak et al., 2019). The asterisk represents the catalytic cysteine required for ubiquitin ligase activity. **D)** Schematic of the *Drosophila melanogaster* CG13025 including the amino acid changes found in the indicated *mus302* alleles; the base substitutions that lead to the amino acid changes are: *D2*, A1T; *Z4933*, G466A; *Z6004*, C400T; *Z1882*, T576A; *D1*/*D3*, T908A. **E)** Survival of heteroallelic *msu302* mutants with a transgene of *CG13025* integrated into *3R* (99F8) (two independent integrants are shown, A and B). Each dot represents a vial, horizontal bar represents the mean and error bars the standard deviation. Horizontal dashed line at Y=1 indicates 100% survival.

*mus302* had been mapped previously between the phenotypic markers *scarlet* (*st*, recombination map 3-44) and *curled* (*cu*, recombination map 3-50) in the third chromosome of *D. melanogaster* (Fig. 1b) (Boyd et al., 1981). This region spans more than 5 Mb and hundreds of predicted genes, so we used recombination mapping to more finely localize *mus302*. Our data showed that *mus302* is close to *st*. We next used deficiency mapping and found that *mus302* is included in a set of 22 genes within the overlap between the deletions *Df(3L)ED4606* and *Df(3L)ED4674*. Analyzing the current literature on the proteins encoded by the genes in this region suggested the predicted gene *CG13025*, which encodes the ortholog of the human RING Finger and WD domain protein 3 (RFWD3), as our primary candidate to be *mus302*. Similar to human RFWD3, CG13025 has an N-terminal RING finger domain (containing the catalytic cysteine), a coiled coil structural motif, and a C-terminal WD domain (Fig. 1c).

We sequenced the *CG13025* coding region of the six *mus302* alleles that were available to us (*D1, D2, D3, Z1882, Z4933* and *Z6004*) and found non-synonymous mutations in all of them that are either nonsense (*Z1882*) or missense mutations (Fig. 1d). *D1* and *D3* had the same mutations, suggesting they originated from the same mutational event or that perhaps stocks were mixed up in the ∼30 years since these mutations were first isolated. Most missense mutations change highly conserved amino acids and are likely to be detrimental to the protein stability or function (*D1, D3, Z4933* and *Z6004*, Fig. S1); the *D2* mutation alters the AUG star codon. Based on the DNA changes and the finding that all mutants are extremely sensitive to a dose of 0.025% MMS (Fig. S2), we conclude that all alleles we analyzed are amorphic or severely hypomorphic.

If the mutant alleles of *mus302* correspond to mutations in *CG13025*, introducing a wild-type copy of *CG13025* should rescue the sensitivity of *mus302* mutants to MMS. We amplified the coding sequence of *CG13025* plus one kb upstream and integrated it into the right arm of the third chromosome (99F8 site) of *D. melanogaster*. Two independent integrants were isolated and recombined onto a chromosome containing the *mus302*^*D1*^ mutation. Flies with this chromosome in *trans* to *mus302*^*Z1882*^ were resistant to 0.05% MMS (Fig. 1e).

The findings that a wild-type copy of *CG13025* rescues the MMS-sensitivity phenotype of *mus302* mutants, and that we found detrimental mutations in all six alleles of *mus302* sequence leads us to conclude that *mus302* is *CG13025* and encodes the *Drosophila* ortholog of *RFWD3*.

### *mus302* is not required for homologous recombination

Human RFWD3 participates in the repair of collapsed replication forks and DSBs through homologous recombination (HR) by ubiquitylating RPA and RAD51, both of which promote HR (Elia et al., 2015; Inano et al., 2017). We hypothesized that *mus302* would work in a similar manner, especially since other the same screen identified other HR genes, including *mus301* (ortholog of *HELQ*) (McCaffrey, St Johnston, & González-Reyes, 2006), and *mus309* (ortholog of *BLM*) (Kusano, Johnson-Schlitz, & Engels, 2001). We tested the sensitivity of *mus302* mutants to hydroxyurea (HU), which stalls replication, and ionizing radiation (IR), which generates DSBs. Surprisingly, *mus302* heteroallelic mutants were not more sensitive to a moderate dose of HU (100 mM) or IR (1000 rads) than their untreated siblings (Fig. 2a, b). Since flies harboring mutations in *spn-A* (encodes the *Drosophila* ortholog of RAD51,), which is required for HR, are sensitive to lower doses of both agents (Brough et al., 2008; Staeva-Vieira et al., 2003), we conclude that Mus302 is not essential for HR.

**Figure 2.**
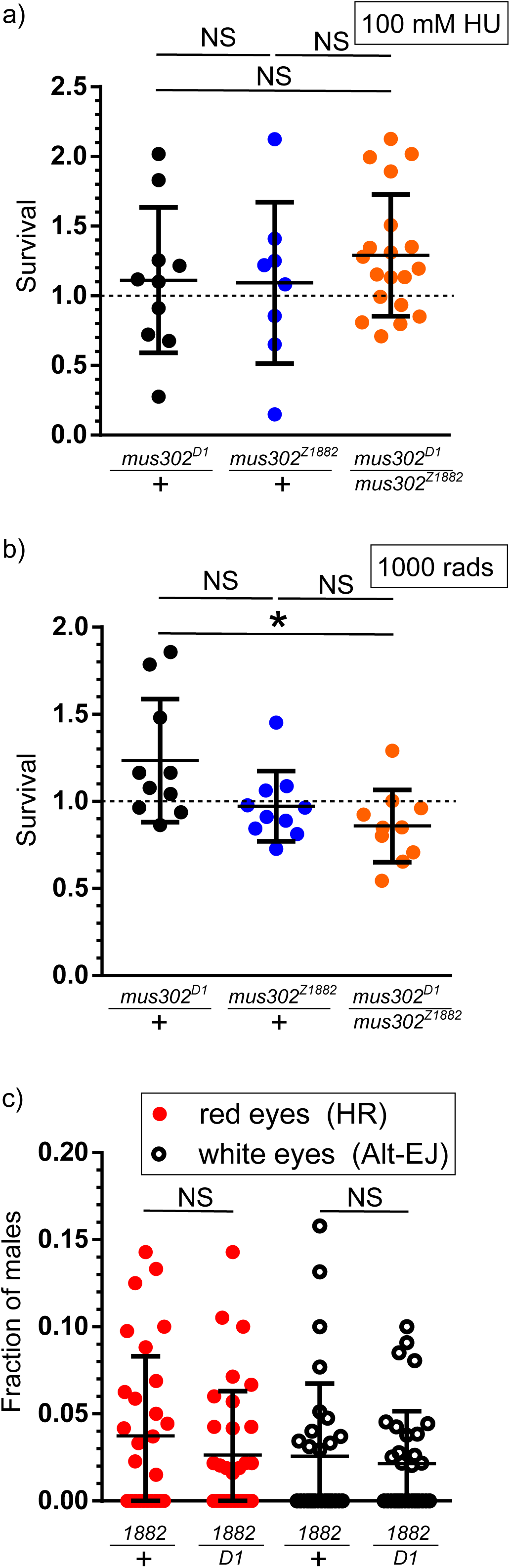
Mus302 is not involved in DSB repair. **A)** and **B)** Survival after exposure to 100 mM hydroxyurea (HU) **(A)** or 1000 rads of ionizing radiation (IR) **(B)**, calculated as in Figure 1A. Horizontal bar represents the mean and error bars the standard deviation. Statistical significance was determined by ANOVA with Bonferroni correction for multiple comparisons (NS, not significant; *p<0.05). Horizontal dashed line at Y=1 indicates 100% survival. **C)** Single males expressing a transposase, containing the *P{wa} P* element, *mus302*^*Z1882*^, and either *mus302*^*D1*^ or not (+) were crossed to females with a compound X chromosome. Each dot represents the fraction of males with either red eyes or white eyes, and not carrying the transposase, per vial. Horizontal bar represents the mean and error bars the standard deviation. Statistical significance was determined by two-tailed t-test (NS, not significant; \**p*<0.05).

In human cells lacking RFWD3, HR repair at either Ter-stalled replication forks or *I-Sce*I-generated DSBs is significantly decreased, as measured by a DR-GFP assay (Elia et al., 2015). We tested the ability of *mus302* deficient flies to perform HR in another type of chromosomal break with a gap repair assay (*P*{*w*^*a*^}) (Adams et al., 2003). This assay takes advantage of a *P* element containing a hypomorphic version of the white gene that confers an orange eye color, inserted in the *X* chromosome. Excision of the *P* element creates a DSB that gives flies a red eye color if repaired by SDSA/HR, or a white eye color when repaired by Polymerase Theta-Mediated End Joining (TMEJ). *mus302* mutant flies exhibit no apparent defect in either repair pathway (Fig 2c).

In contrast to cells deficient in RFWD3, *mus302* mutants are not sensitive to HU or IR and are proficient in SDSA. We conclude that the functions described for human RFWD3 are not shared with the *Drosophila* ortholog.

### Mus302 functions independently of Spn-A

Given that both *mus302* and *spn-A* (the *RAD51* ortholog) mutants are sensitive to MMS (albeit different MMS concentrations are required to see such sensitivity (Staeva-Vieira et al., 2003)) and that RAD51 has functions outside of HR, it remains formally possible that they are part of the same pathway. Hence, we directly tested such possibility.

We exposed *mus302* and *spn-A* single and double mutants to increasing concentrations of MMS (0%, 0.001%, 0.025%). In untreated flies, we did not observe any differences in viability between the three genotypes (Fig. 3); however, at the low dose of 0.001% MMS, *mus302* s*pn-A* double mutants had significantly reduced survival compared to *mus302* single mutants (Fig. 3). As previously reported, a dose of 0.025% MMS is lethal for *mus302* single mutants but not for *spn-A* mutants (Boyd et al., 1981; Staeva-Vieira et al., 2003); double mutants are also highly sensitive to this dose (Fig. 3). These results show that, unlike their human orthologs, Mus302 and Spn-A are part of different DNA repair pathways.

**Figure 3.**
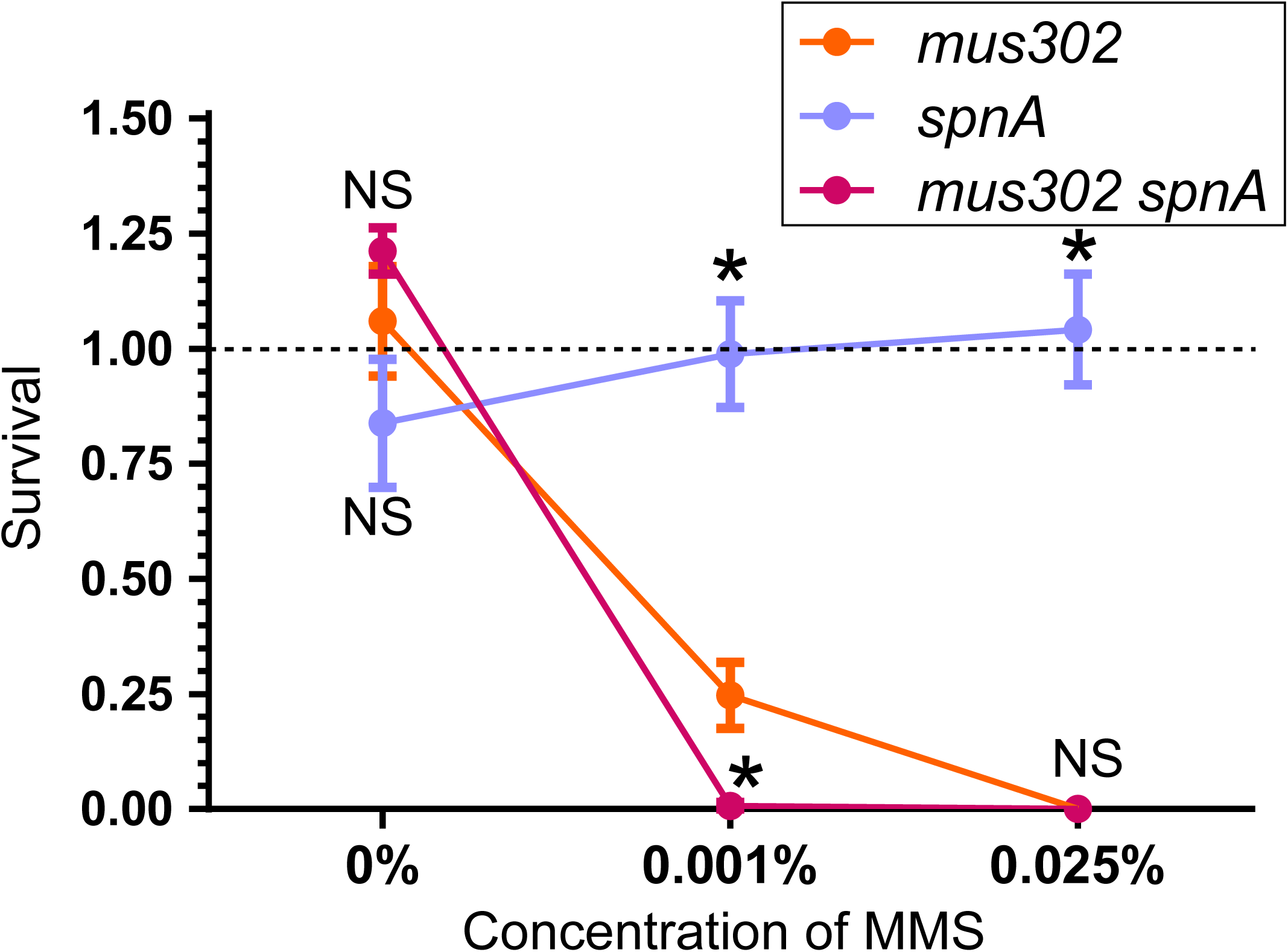
Mus302 functions independently of Spn-A. Survival after exposure to the indicated dose of MMS was calculated as in Fig. 1A. Dots represent the mean and error bars the standard error of the mean (*n* ≥ 5 biological replicates). Statistical significance was determined by ANOVA with Bonferroni correction for multiple comparisons (NS, not significant; \**p*<0.05) for each concentration of MMS. An outlier was removed from the *spn-A*, 0.001% MMS with ROUT test, Q = 1%. Horizontal dashed line at Y=1 indicates 100% survival.

### ATR phosphorylation motifs of RFWD3 appeared late in evolution

To understand the functional differences observed between the human and the *Drosophila* orthologs, we performed a protein sequence alignment between different RFWD3 orthologs. In addition to the human and the fly proteins, we used sequences from five other animal species: mouse (*Mus musculus*), chicken (*Gallus gallus*), frog (*Xenopus tropicalis*), zebrafish (*Danio rerio*) and sea urchin (*Strongylocentrotus purpuratus*). We observed a high conservation across species from the beginning of the RING finger through the end of the protein. However, the sequence upstream of the RING finger showed low conservation (Fig. 4).

**Figure 4.**
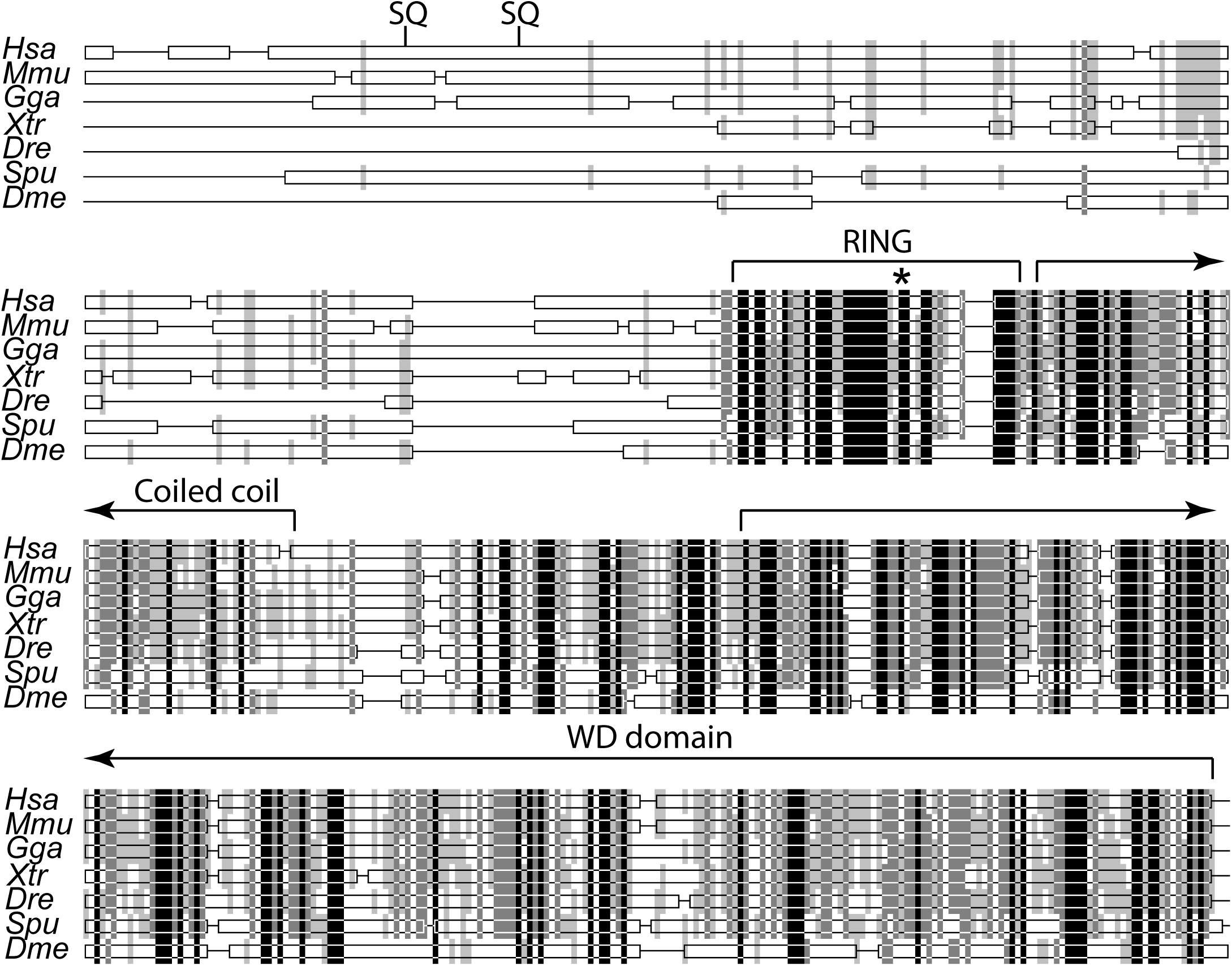
The N terminus of RFWD3 appeared late in evolution. Protein alignment of 7 RFWD3 orthologs performed as in Figure S2. Thin lines represent gaps introduced for optimal alignment. Black bars indicate conservation across all species examined; light colors represent conservation in a subset of species (see Fig S2). SQ indicates the two SQ motifs in human RFWD3 known to be phosphorylated by ATR. Domain boundaries shown for the human protein determined as in Figure 1c. Asterisk indicates the catalytic cysteine.

Since it is the C-terminus of the human RFWD3 that interacts with RPA32 (Liu et al., 2011), we reasoned that this interaction may be conserved. Moreover, four amino acids in RPA32 required for its interaction with RFWD3 are present in flies (Feeney et al., 2017). In contrast, the N-terminus of the human protein has two serines (S46 and S63) that are part of SQ motifs that are phosphorylated by ATR in response to DNA damage (Fu et al., 2010). They are also hypothesized to target RFWD3 repair to S phase. Strikingly, we observed that both serines are missing in the frog, zebrafish, and fly orthologs, and at least one is missing in the chicken and sea urchin proteins (there is a nearby SQ motif in these latter two species but the surrounding amino acids sequences are not conserved).

Based on our analysis, we suggest that ATR phosphorylation of RFWD3 was acquired relatively recently on the mammalian branch. We speculate that Mus302 and other non-mammalian orthologs may be active outside of S phase, and that this may represent the ancestral function of the protein. This would explain our observation that Mus302 is not involved in homologous recombination, a DNA repair pathway most active during S phase in some organisms.

Mus302 is required for survival in the presence of MMS. Alkylating damage is repaired outside of S phase by excision repair mechanisms (Kondo, Takahashi, Ono, & Ohnishi, 2010). Because most of the protein sequence of RFWD3 is conserved, it is possible that the human protein is also involved in the repair of alkylating damage outside of S phase, and that *mus302* represents a “separation-of-function” ortholog that can be used to elucidate possible functions of RFWD3 in excision repair pathways.

In summary, we have found that the mutagen sensitive complementation group *mus302* corresponds to the *Drosophila melanogaster* ortholog of the human *RFWD3*. The findings presented here show that Mus302 lacks the known functions of RFWD3 in promoting homologous recombination during replication fork collapse and DSB repair. Our analysis suggests that Mus302 may not be phosphorylated by the ATM/ATR kinases and we propose that this is responsible for the differences between the fly and the human protein. Further characterization of this gene in *Drosophila* has the potential to uncover new functions of the human protein.

## Acknowledgements

We thank Evan Dewey for comments on the manuscript. This work was supported by a grant from the National Institute of General Medical Sciences to J.S. under award 1R35GM118127 and by a grant from the National Cancer Institute to D.R. under award 5R01CA222092.

## Figure legends

**Figure S1.** Representation of an alignment between The RFWD3 protein sequences from *Homo sapiens* (*Has*), *Mus musculus* (*Mmu*), *Gallus gallus* (*Gga*), *Xenopus tropicalis* (*Xtr*), *Danio rerio* (*Dre*), *Strongylocentrotus purpuratus* (*Spu*), and *Drosophila melanogaster* (*Dme*) were obtained from Ensembl and aligned with ClustalX. Alignment representation was made with GeneDoc. The colors represent the conservation of the identity of the amino acid, (four levels of conservation: black, present in all seven species, dark grey, in 6/7, light grey, in 4/7 or 5/7, and white, in <4/7); hyphens indicate gaps introduced for optimal alignment. Arrows denote the residues mutated *mus302* alleles, with the predicted new amino acid shown below.

**Figure S2.** Survival after exposure to 0.025% MMS calculated as in Fig. 1A. Mutants were hemizygous for the indicated *mus302* allele over *Df(3L)ED4606*. Each dot represents a vial, horizontal bar represents the mean and error bars the standard deviation.

